# High-throughput cell-free profiling of SARS-CoV-2 RBD variants enables rapid and quantitative in vitro affinity landscape mapping

**DOI:** 10.1101/2025.09.23.678000

**Authors:** Laura Grasemann, Jiami Han, Julia Tischler, Fatemeh Arefi, Maria Andrea Gonzales Castillo, Edward B Irvine, Ningning Chen, Sai T. Reddy, Sebastian J. Maerkl

## Abstract

SARS-CoV-2 variants continue to threaten public health, necessitating the study of cumulative and epistatic effects of receptor-binding domain (RBD) mutations on antibody evasion. We present a high-throughput platform combining cell-free protein synthesis and microfluidics to quantify the affinity of a large number of RBD triplet mutants covering the evolutionary space between wild-type and Omicron against two therapeutic antibodies and one engineered binder. Using rapid in vitro gene assembly and cell-free synthesis, we expressed 518 RBD variants and obtained 31,740 quantitative affinity measurements to generate three comprehensive binding energy landscapes. This approach enables rapid and large-scale in vitro affinity profiling and machine learning-based predictions, providing a valuable tool for studying emerging variants.

## Main

The rapid accumulation of mutations in the receptor-binding domain (RBD) of the SARS-CoV-2 spike protein has led to significant immune escape from neutralizing antibodies ^1–3^. Two prime examples of these are the therapeutic antibodies Bamlanivimab (LY-CoV555) and Imdevimab (REGN10987) ^4,5^. While single mutations are well studied, the combinatorial effects of multiple mutations remain challenging due to the vast sequence space and complex residue interactions ^6^. Binding involves multiple interacting residues stabilized by others, making isolated mutations insufficient for understanding escape mechanisms. Several studies have leveraged high-throughput assays and machine learning to map the functional consequences of combinatorial RBD mutations, revealing how concurrent substitutions can affect antibody binding and viral fitness ^7–9^. However, mapping mutational landscapes to quantitative affinity measurements, rather than binary binding outcomes, remains constrained by the limited throughput of quantitative affinity assays such as surface plasmon resonance (SPR) and biolayer interferometry (BLI).

Advancements in microfluidics and cell-free protein expression have enabled high-throughput protein-ligand affinity measurements ^10^. Cell-free systems bypass the need for living cells for protein synthesis ^11,12^.

Microfluidic platforms enable high-throughput quantitative molecular interaction measurements ^13^. Combined, these technologies facilitate rapid and large-scale characterization of protein-protein interactions ^14^.

Here, we present a platform combining cell-free protein synthesis and microfluidics for high-throughput binding affinity measurements of SARS-CoV-2 RBD and neutralizing antibodies. We generate affinity data for combinatorial triple-mutation variants of the RBD to train machine learning models that predict affinity from RBD protein sequences. Our approach enables rapid assessment of combinatorial mutations and improves sequence-to-affinity-value modeling, more precisely than binary label modeling, offering a valuable tool for protein engineering and antibody design, with potential applications in developing therapeutics against evolving pathogens like SARS-CoV-2.

We designed a combinatorial RBD library of 560 variants, each containing three mutations selected from the 16 mutations present in Omicron BA.1 relative to the ancestral RBD of SARS-CoV-2 (Wu-Hu-1) (Fig. 1A-C). This choice mimics spontaneous mutational events that may occur without selection pressure, providing insight into affinity changes occurring at the early stages of viral evolution toward immune escape. The library was synthesized using a segment-based assembly approach (Fig. 1D) dividing the RBD gene into 6 segments 126– 283 bp in length, requiring only a total of 50 different gene blocks to synthesize all 560 variants, which markedly reduced per-variant gene synthesis cost. We successfully assembled 518 variants using this approach, which corresponds to a 92.6% success rate. The resulting PCR product was directly used for cell-free expression in the PURE system supplemented with a disulfide bond enhancer, ensuring efficient expression and proper folding of RBD variants. The expressed variants were then diluted and directly spotted onto a glass slide without purification using an automated microarray spotter and aligned to a microfluidic chip (Fig. 1E). All variants were expressed as RBD-GFP fusion proteins and immobilized on-chip via GFP binding to a surface immobilized anti-GFP antibody, ensuring 1:1 stoichiometry of GFP: RBD for normalized binding measurements (Fig. 1G, H). Using the MITOMI approach ^13^, we measured the binding affinities of two antibodies previously used as therapeutics, Bamlanivimab (Eli Lilly, LY-CoV555) and Imdevimab (Regeneron, REGN10987), due to their distinct neutralisation profiles against variants of concern, and a novel binder DBR3_03 ^15^, as it has computationally designed broad-spectrum efficacy (Fig. 1C). We quantified binding events by antibody fluorescent detection at five antibody concentrations sequentially, then determined relative affinities by linear regression (Fig. 1I). Relative affinity values were inferred from intensity values using their quasi-linear correlation at low ligand concentrations ^16^, bypassing the need for saturation binding curves (Fig. 1H). Chip-to-chip reproducibility was confirmed with Imdevimab measurements (R^2^ = 0.89, Fig. S4).

**Fig. 1.**
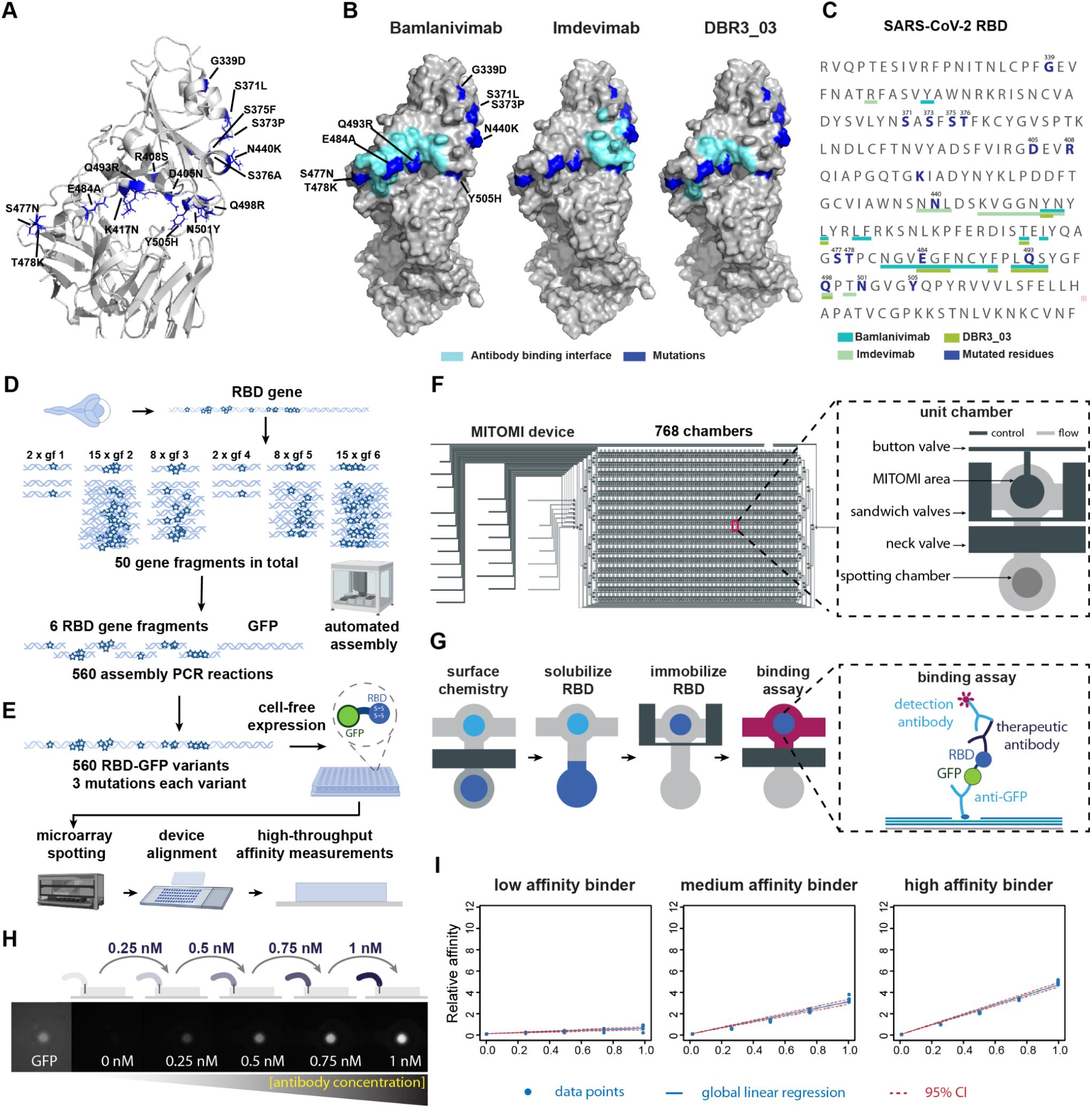
Design of RBD triple mutation library, cell-free variant expression and microfluidic affinity assay. (A) Ribbon model of the wild-type SARS CoV-19 spike protein RBD. Residues that are mutated in omicron are highlighted in dark blue. (B) Surface model of the wild-type SARS CoV-19 spike protein RBD. Respective binding interfaces for Bamlanivimab, Imdevimab, and DBR3_03 are highlighted in light blue. Residues that are affected by omicron mutations are highlighted in dark blue. (C) Amino acid sequence of wild-type RBD. Residues that are mutated in omicron are highlighted in dark blue. Residues involved in the binding surface of Bamlanivimab, Imdevimab, and DBR3_03 are underlined in shades of green. (D) Workflow for RBD library construction. (E) Cell-free expression and high-throughput affinity measurement workflow. (F) MITOMI device schematic: Two-layer chip with 768 unit chambers, featuring valves for chamber isolation and reagent control. (G) On-chip assay overview: RBDs are immobilized in individual chambers, followed by a fluorescence-based binding assay with therapeutic antibodies or binders. (H) Binding assay workflow: GFP signal is measured for expression level, then antibody titrations (0–1 nM) and detection antibody are applied. (I) Example binding curves for Imdevimab: Low (Omicron), medium (variant 170), and high (wild type) affinity binders, with regression fits and confidence intervals.

Our MITOMI system (Fig. 1F) with 768 reaction chambers enables high-throughput and quantitative affinity measurements over a broad range of affinities. Its capacity allows for the measurement of all 518 variants in four replicates, requiring only three chips per antibody. Across all three binders, this approach generated 31,740 affinity measurements, demonstrating its capability of obtaining large-scale binding data. Details are provided in the Supplementary Information (Supplementary Fig. 1).

We converted the measured affinities to normalised relative affinities (Fig. 2A) and defined “mean relative affinity” to quantify the contribution of single mutations in the context of triple mutations (Fig. 2B). Bamlanivimab binding was mainly affected by RBD mutations E484A and Q493R (present in antibody-binding epitope) and enhanced by S371L (outside of epitope, possibly via increased hydrogen bonding) ^17^. For Imdevimab, both epitope and non-epitope mutations reduced affinity. DBR3_03 showed robustness to RBD epitope mutations but was impacted by S375F, T376A, and D405N. Notably, S375F and T376A reduced affinity for both Imdevimab and DBR3_03 (∼0.5-fold and ∼0.6-fold), suggesting they may destabilize a shared binding region.

**Fig. 2.**
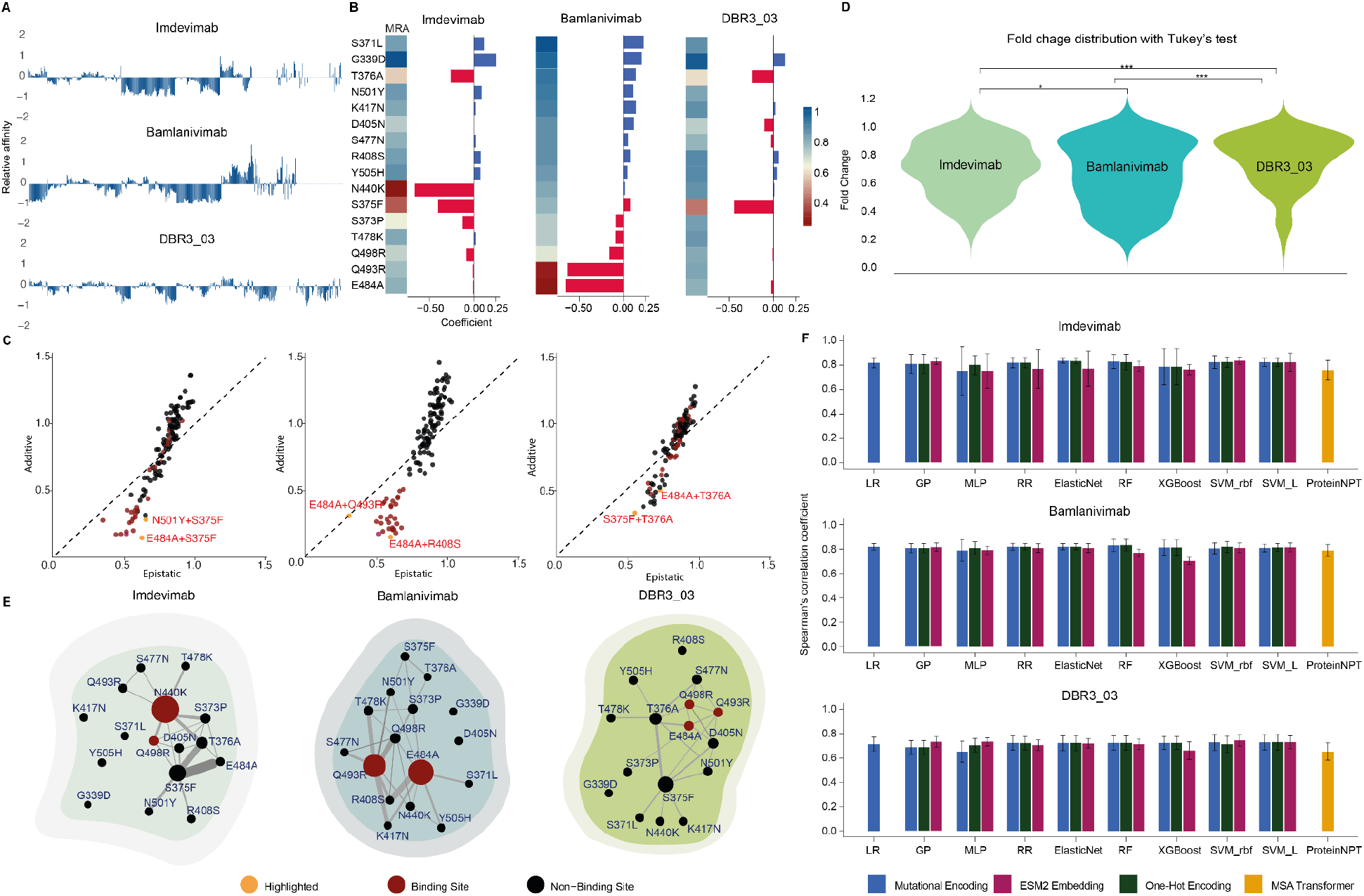
Analysis of mutational synergies and sequence-based affinity prediction benchmarking. (A) Data overview of three antibodies. Each bar represents one RBD variant, and the variant order is identical across the three panels. Values are normalized to WT (WT = 1) and mean-centered (WT → 0). (B) The heatmap on the left depicts mean relative affinity (MRA) caused by single mutations in the three datasets for each binder tested, red represents a stronger affinity-reducing effect. The bar plot on the right shows the ElasticNet coefficient of all mutation flags, with negative values representing negative impact on affinity-reducing effect. (C) Synergy on binding affinity (y) vs. additive effect of double-mutation pairs (x), dashed line gives reference for x=y, yellow dots are highlighted high synergy pairs, variants in red contain at least one mutation inside the epitope. Variants in yellow are annotated with mutations. (D) Distributions of binding affinity shifts across antibodies. Violin plots show the fold-change (only fold-change < 1 included) for triple mutation variants. DBR3_03 exhibits the highest median affinity with the lowest dispersion, indicating superior resistance to escape mutations. (E) Mutational network mapping of synergistic effect between escape mutations. Thickness of the edge between nodes is proportional to the additional affinity-reducing effect caused by the connected pairs of mutations (only fold-change < 1 connected), while the size of nodes is proportional to the affinity-reducing effect of an individual mutation. Nodes in red are located inside the epitope. (F) Spearman correlation (mean ± SD, n = 5 splits) between predicted and measured affinities, trained with four sequence representations. Results are displayed separately for each binder.

To highlight key synergistic interactions driving antibody escape, we plotted the mean relative affinity of every double mutant against the additive expectation from its two single mutants (Fig. 2C). Deviations from the 45° line quantify epistasis. For Imdevimab, E484A + S375F exhibits strong synergy, despite E484A alone not being a top escape mutation, suggesting combinatorial effects drive immune resistance. Similarly, N501Y, which has a minor impact alone, amplifies affinity loss when paired with S375F, underscoring the context-dependent nature of escape mutations. For Bamlanivimab, E484A and Q493R independently reduce affinity but show no additional effect together, indicating distinct escape mechanisms. In contrast, R408S, a weaker single mutation, exhibits the strongest affinity reduction when paired with E484A, suggesting a destabilizing interaction. The fact that E484A synergizes with different partners across antibodies aligns with its prevalence in circulating variants. Notably, Bamlanivimab’s most potent synergies occur in its RBD epitope. Mutation pairs exhibiting a strong synergistic affinity-reducing effect are distinctly clustered below the diagonal line, whereas Imdevimab and DBR3_03 exhibit strong combinatorial effects from mutations outside their epitopes, revealing distinct vulnerability patterns. Binding affinity fold change distributions (Fig. 2D) highlight the robustness of DBR3_03. The violin plot shows that DBR3_03 has the highest mean affinity and lowest variability, indicating potentially reduced susceptibility to escape mutations (Fig. 2D). Tukey’s test confirms DBR3_03’s significantly higher affinity retention compared to Bamlanivimab (p < 0.001) and Imdevimab (p < 0.001), while Imdevimab and Bamlanivimab exhibit comparable reductions (p = 0.997). Structural factors likely contribute to DBR3_03’s mutational resilience. Unlike Bamlanivimab and Imdevimab, which rely on binding to epitope surfaces with flexible loops in CDRs ^18^, DBR3_03 binds via a compact, two-helix scaffold, engaging the RBD with a mix of backbone and side-chain interactions rather than extensive side-chain contacts alone ^15^. This reduces its dependence on mutable RBD residues, explaining its higher tolerance to escape mutations.

To further dissect antibody escape vulnerabilities, we constructed mutational networks (Fig. 2E) to reveal escape hotspots. DBR3_03, despite its overall resistance, exhibits an escape hotspot centered around S375F and T376A, suggesting a potential weak point for future mutational escape. While Bamlanivimab’s escape profile is dominated by E484A and Q493R, with additional destabilizing interactions from K417N, R408S, and Q498R, Imdevimab is more susceptible to mutations outside its epitope, particularly N440K and the synergistic S375F + E484A pair, which likely disrupts structural integrity. These findings provide valuable guidance for studying immune escape mechanisms and inform the rational design of more resilient binders that minimize immune escape risks.

Higher-order mutational synergy was assessed by comparing triple mutations based on fold change (Supplementary Data). For example, S477N, T478K, and Q498R dramatically reduce Bamlanivimab affinity (fold change = 0.11), a far greater reduction than any double combination of these mutations. This suggests a non-additive interaction amplifying escape, reinforcing the need for combinatorial analyses to capture mutational effects that would be missed in single point mutation studies.

To test whether these univariate trends persist after conditioning on all other mutations, we fitted an Elastic-Net to the 16 binary flags. Elastic-Net blends L^1^ (lasso) and L^2^ (ridge) penalties, where it both zeros out negligible weights, leaving a sparse set of drivers, and shrinks correlated coefficients toward each other for stability. This yields a robust, interpretable estimate of each residue’s intrinsic effect on affinity, conditioned on the other 15 mutations. The negative coefficients of each antibody match the residues that have a negative effect on binding in mean affinity analysis (Fig. 2B), demonstrating that their effects are intrinsic rather than artefacts of library composition. Because Elastic-Net attains a decent accuracy while keeping its coefficients directly interpretable as per-mutation affinity changes, we envision it as a mechanistic compass to highlight escape drivers, then proceed to examine higher-capacity models for maximising predictive power.

We next benchmarked three different RBD sequence representations and eight machine learning models for their performance in predicting relative affinity. To encode the sequences, we used three approaches: mutational encoding, one-hot encoding, and a pre-trained protein language model embedding by ESM2 ^19^. In mutational encoding, each possible mutation was assigned a binary value (1 for presence, 0 for absence), creating a vector that tracks differences relative to the ancestral RBD sequence (Wu-Hu-1). One-hot encoding retains full sequence information, while ESM2 embeddings can capture structural and evolutionary information. Mutational and one-hot encoding provide sparse, interpretable feature spaces that emphasize explicit sequence differences, whereas ESM2 embeddings offer a dense, high-dimensional representation that enhances the ability to model nonlinear relationships but complicates feature interpretation and distance-based learning. Each method has distinct advantages: simpler encodings benefit models that rely on feature separability, while ESM2 embeddings provide richer biological context for models capable of capturing complex patterns.

These embeddings served as input features for machine learning models that included Gaussian Process (GP), Linear Regression, Ridge Regression, XGBoost, Support Vector Machine (SVM), and Multi-Layer Perceptron (MLP), Elastic Net, and Random Forest. Model performance was assessed using Pearson and Spearman correlation coefficients, mean squared error (MSE), and R^2^.

Across all antibodies, the top-performing models achieved Spearman correlations ranging from 0.75 to 0.86 in random-split five-fold cross-validation, indicating robust sequence-affinity relationships and dataset suitability for modeling (Fig. 2F). Optimal embeddings and model performance were dataset-dependent (Supplementary Table 1–3). Notably, the Imdevimab dataset, despite its greater complexity due to synergistic effects and smaller size (18% fewer data points than DBR3_03), showed no performance degradation. While ESM2 embeddings encode rich biological information, their high-dimensional complexity may introduce noise, given limited data size.

We next tested ProteinNPT, a transformer-based model that leverages sequence, MSA, and label information for protein property prediction ^20^. It models sequence context with a residue-to-residue self-attention mechanism, allowing the model to capture epistatic effects without interaction terms. We trained ProteinNPT models with supervised affinity prediction and MSA Transformer embeddings plus auxiliary zero-shot fitness labels. The result achieved Spearman correlations of 0.79, 0.76 and 0.65, for Bamlanivimab, Imdevimab and DBR3_03, respectively (Fig. 2F). These scores were comparable to, but did not exceed, the best-performing baseline models, suggesting that performance gains require more data and greater alignment of the task and the pretrained model.

By combining in vitro gene assembly with cell-free protein synthesis and high-throughput microfluidic molecular interaction analysis we were able to rapidly establish three comprehensive and quantitative molecular binding landscapes, mapping the escape potential of SARS-CoV-2 RBD variants. This completely in vitro approach enables the rapid generation and quantification of protein binders on a large scale (hundreds of protein variants and tens of thousands of measurements). Overall, as more data becomes available through this high-throughput pipeline, sequence-based affinity prediction can be further refined, improving our understanding of how mutations influence molecular interactions. Expanding these datasets will enhance predictive accuracy and support the development of more efficient computational models for antibody affinity estimation. More generally, this approach enables near-immediate evaluation of new mutations and their effect on antibody binding, which is essential for real-time surveillance, predicting antibody-escape potential, and guiding therapeutic antibody design.

## Methods

### Variant Library Generation

The sequence for the RBD is based on a previous publication^21^ and was codon-optimized for *E. coli* using the GenScript codon optimization tool. The DNA sequence for the wild type and Omicron can be found in the Supplementary Information. Mutations were incorporated into the sequence by manually mutating the respective amino acids while changing the overall sequence as little as possible, ensuring that the codon usage is optimal for *E. coli*. Reference mutations for Omicron dated from September 2022 and were taken from outbreak.info. Mutations were selected based on those with an occurrence of at least 50% by September 2022 ^22^. These include 16 mutations: G339D, S371L, S373P, S375F, T376A, D405N, R408S, K417N, N440K, S477N, T478K, E484A, Q493R, Q498R, N501Y, and Y505H.

The gene for the RBD-GFP fusion protein was divided into six fragments (g1-g6) (Fig. 1D) for the RBD section and one fragment for GFP, all with overlaps of 22–31 bp and similar melting temperatures for said overlaps.

The six fragments incorporated all possible mutations between the wild type and Omicron. A schematic of the gene and the g-block design, including mutation distribution, is shown in Fig. S3. The spread of the mutations across the six g-blocks included between 1 to 4 mutations, leading to 2^n^ versions of each g-block, with n being the number of mutations in the g-block. We thus ordered 2 versions of g1, 16 versions of g2, 8 versions of g3, 2 versions of g4, 8 versions of g5, 16 versions of g6, and one version for GFP. The DNA for all g-blocks is listed in the Supplementary data g-block list file. All g-blocks and primers were ordered from IDT.

RBD-GFP variants were created by assembling the respective g-blocks using assembly PCR with Taq polymerase (NEB). 24 variants were assembled at a time. A master mix, including the g-block for GFP but omitting the DNA polymerase and primers, was prepared (1.66x Standard Taq Reaction Buffer, 333 μM dNTPs [Thermo Fisher], ultrapure water, 1.66 nM g-GFP). 9 μL of the reaction mix was added to each well of a 96-well plate (Eppendorf, twin.tec PCR plate). Subsequently, 1 μL of the respective g-blocks 1–6 (1 nM stock solution) was added to the reaction mix, resulting in a total reaction volume of 15 μL. Both steps were performed using an EP Motion 5073 (Eppendorf) Liquid Handling System. The reactions were transferred to PCR strips (VWR), and 0.2 μL of Taq polymerase (5000 U/mL stock solution, NEB) was added. First, a 10-step touchdown PCR was run on a thermocycler, as detailed in Supplementary Information Table 2. During touchdown PCR, the initial annealing temperature is set above the projected melting temperature of the primers or, as in this case, the melting temperature of the overlaps. With every cycle, the annealing temperature decreases. Touchdown PCR increases the sensitivity and specificity of PCR reactions.

After touchdown PCR, B5’final and B3’final (sequences detailed in Section) were added to the reaction mix at a final concentration of 0.2 μM, and a standard Taq PCR was performed, as detailed in Supplementary Information Table 3. The product of this PCR was then transferred to a fresh 50 μL Taq PCR reaction (1x reaction buffer, 200 μM dNTPs, 0.2 μM of each primer, 0.25 μL Taq polymerase, and 0.5 μL reaction product) using the conditions detailed in Supplementary Table 3. The product was verified using gel electrophoresis. About 87% of the 560 designed variants showed a clean PCR product. Another 7% of the variants did not exhibit a clean band during the second PCR step but did show a clean band after the first PCR amplification. These were added to the library as assembly PCR products. In total, we successfully synthesized DNA for 518 variants. The synthesized DNA was used for expression without further purification steps.

### PURE Expression of Variants

The variant library generated in the previous section was divided into three parts. Plate 1 contained variants 1– 184, plate 2 contained variants 185–346, and plate 3 contained variants 347–518. Each plate additionally contained two replicates for the wild type and one to two replicates for Omicron. Variants were expressed as RBD-GFP fusion proteins. The PUREfrex 2.1 system was used with a DsbC supplement to enable disulfide bond formation. Disulfide bonds are critical for the folding of the RBD ^23^. Reactions were set up according to the supplier’s manual for the usage of the DsbC set, with the addition of Protector RNase Inhibitor (Roche). For each 20 μL of reaction, this included 4.5 μL ultrapure H_2_O, 8 μL Solution I, 1 μL of 10 mM Cysteine, 1 μL of 80 mM GSH, 1 μL of 60 mM GSSG, 1 μL of Solution II, 2 μL of Solution III, 1 μL of 80 μM DsbC, and 0.5 μL of RNase inhibitor. One master mix was prepared for each plate. 9 μL of the master mix were distributed to the wells of a conical 384 microwell plate (ArrayIt, MMP384). Subsequently, 0.5 μL of DNA for one variant from the library was added to each well using a multichannel pipette. The plate was sealed with an alumina plate seal foil (Eppendorf) and incubated using a ThermoMixer (Eppendorf) and a ThermoTop (Eppendorf) for 4 h at 37°C while shaking at 400 rpm. The samples were diluted with 20 μL of 2% (w/v) BSA in ultrapure water to increase the total sample volume. The plate was used directly for microarray spotting as detailed in the next section and subsequently stored at −20°C.

### Microfluidic Chip Fabrication

Detailed designs for the flow and control layers of the microfluidic device are available for downloading (lbnc.epfl.ch). The molds for each layer were fabricated on separate wafers using standard photolithography techniques on a 4-inch silicon wafer as previously reported ^24^. SU-8 negative photoresist (GM 1070, Gersteltc SARL) was used to generate the channel features of the control layer at a height of 30 μm. The flow layer was fabricated using AZ 10XT-60 positive photoresist (Microchemicals GmbH) at a height of 15 μm. The flow layer mold was annealed at 180°C for 2 h in a convection oven to round the features of the flow channel. Before its first use, each wafer was pretreated with TMCS (trimethylchlorosilane) overnight. For soft lithography chip fabrication of the control layer, the mold was pretreated with TMCS for 5 min. PDMS with a ratio of elastomer to cross-linker of 5:1 was prepared and poured over the wafers. Subsequently, the PDMS-coated wafer was desiccated for 30 min, followed by a partial curing step of 20 min at 80°C. For the flow layer, a PDMS to cross-linker ratio of 20:1 was used and spin-coated onto the mold for the flow layer at 400 rcf to achieve a height of 50 μm. The PDMS-coated mold was then partially cured at 80°C for 25 min. Subsequently, the devices for the control layer were cut out, and the control layer’s inlets were punched using a precision manual punching device (Syneo, USA). The control layer was then aligned on top of the flow layer, and the microfluidic chip was placed in the oven for 90 min at 80°C to allow the layers to bond and the PDMS to fully cure. The bonded chips were cut and removed from the flow layer mold, and the fluid entry points for the flow layer were punched.

### Microarray Spotting

#### 1. Epoxy-Silane Glass Slide Preparation

Epoxy-silane glass slides for microarray spotting were prepared as previously published ^25,26^, with a slightly adapted protocol. A 25% (v/v) solution of ammonia in MilliQ water was heated to 80°C. Subsequently, 30% (v/v) H_2_O_2_ was added to the mixture, and the glass slides were incubated in the mixture for 30 min. The glass slides were then washed in MilliQ water and dried with N_2_. Next, the glass slides were placed in a bath containing 0.62% (v/v) 3-glycidoxypropyltrimethoxysilane (3-GPS) in toluene for 20 min at room temperature. The glass slides were washed with fresh toluene to remove any unbound 3-GPS and baked at 80°C for 60 min. The glass slides were stored in opaque boxes under vacuum until further use. Prior to sample microarray spotting, the glass slides were rinsed with toluene and dried with N_2_.

#### 2. Sample Microarray Spotting

The variants were spotted onto an epoxy-coated glass slide using a QArray2 microarrayer (Genetix) with an MP3 microarray printing pin (ArrayIt). Four replicates were spotted per variant using a randomized spotting pattern, with two stamps per spot (stamping time: 50 ms; inking time: 100 ms). Subsequently, the PDMS chip was aligned on top of the spots and bonded overnight at 50°C. This temperature ensured sufficient bonding strength while minimizing protein degradation. Six chips were fabricated, spotted, and bonded on the same day as the PURE expression of the respective variants. After overnight bonding, the remaining chips were stored under vacuum until further use.

### On-Chip Binding Assays

#### 1. Surface Derivatization and Immobilization of RBD-GFP Variants

Control lines were filled with MilliQ water and pressurized with air at 10 psi until all lines were filled. Subsequently, the pressure was gradually increased to 22 psi. The pressure for the flow lines was set to 4 psi unless specified otherwise. The spotting chambers were isolated by closing the neck valve at all times unless specified otherwise. The surface treatment was automated using a custom Python script and involved the following steps, as illustrated in Supplementary Fig. 1:

1. The chip was primed with 2 mg/mL Biotin-BSA in PBS (Thermo Fisher, 29130) for 20 min.
2. NeutrAvidin (Thermo Fisher, 3100) was flown at a concentration of 1 mg/mL in PBS for 20 min.
3. Button valves were actuated, and Biotin-BSA was flown for another 40 min to block all NeutrAvidin binding sites except those under the button area.
4. Washing with 0.005% (v/v) Tween 20 (Sigma, P1379) in 0.5 M PBS was performed for 5 min after each step to remove unbound components.

Next, 20 μg/mL of biotinylated anti-GFP antibody (Abcam, ab6658) in 2% (w/v) BSA and 1 M PBS, supplemented with 25% chicken serum (Sigma, C5405), was flown for 2 min with buttons closed and for 15 min with buttons open to allow the antibody to bind to the NeutrAvidin residues exposed under the button area. Chicken serum was used to passivate the surface and prevent nonspecific binding. Subsequently, buttons were actuated, and unbound components were washed away for 5 min with Biotin-BSA solution.

After solubilization, the RBD-GFP variants were immobilized by incubating for 1 h with closed buttons to equilibrate concentrations and for another hour with open buttons to allow binding to anti-GFP antibodies. GFP fluorescence was measured using a FITC filter to determine the amount of bound RBD-GFP.

#### 2. Binding Assay with Therapeutic Antibodies

Binding assays were conducted with Bamlanivimab (ProteoGenix, PX-TA1031), Imdevimab (ProteoGenix, PTX-COV-A553), and the previously published binder DBR3_03 ^15^. DBR3_03 included an Fc-tag and was kindly provided by Bruno Correia and Anthony Marchand.

Antibody solutions at different concentrations (0 nM, 0.25 nM, 0.5 nM, 0.75 nM, and 1 nM) were prepared in 2% (w/v) BSA and 1 M PBS, supplemented with 25% chicken serum. For each concentration, the following steps were repeated:

1. Antibody solution was flown for 2 min with buttons actuated and for 13 min with buttons open.
2. The flow was stopped, and the solution was left to equilibrate for 7 min.
3. Buttons were actuated, and the chip was washed with PBS-Tween for 5 min.
4. Detection antibody (Goat Anti-Human IgG Fc [SureLight PE], Abcam, ab131612) diluted to 2 μg/mL in 2% (w/v) BSA and 1 M PBS was flown for 2 min with buttons closed and for 5 min with buttons open.
5. The chip was washed again with PBS-Tween for 5 min.

Fluorescence was imaged using a Nikon ECLIPSE Ti microscope with a Cy3 filter and an exposure time of 200 ms.

### Image Processing

Images were processed using a custom script based on FIJI, as described previously ^27^. A circular region corresponding to the MITOMI valve area was detected based on the brightfield image using the Hough Circle Transform plugin (UCB Vision Sciences library). After 30 steps of erosion, a region of interest (ROI) was defined as the button area. The MITOMI valve area was then dilated by 30 steps to generate the ROI for the background detection. To ensure the mask was well-aligned with the signal image, a stitch of both images was merged. If the mask did not align with the signal of the fluorescent image, the respective images were measured manually using FIJI’s built-in measuring tool applied to a button-sized ROI.

### Data Analysis

For each unit cell, the median fluorescence value of the background area was subtracted from the median fluorescence value of the button area to calculate background-subtracted fluorescence. A threshold of 500 relative fluorescence units (RFU) was applied to background-subtracted FITC values to exclude spots with insufficient RBD-GFP binding. The Cy3 channel’s median background-subtracted value was normalized by dividing it by the corresponding FITC channel value to account for the amount of pulled-down RBD-GFP.

Fluorescence values at 0 nM antibody concentration were subtracted from all other concentrations to remove nonspecific signals. The resulting values were plotted against antibody concentrations, and a linear regression model (y = ax) was fitted using Python’s LinearRegression function from sklearn.linear_model. The slope of the regression (a), representing relative affinity, was calculated. The standard error of the slope (SES) was estimated, and confidence intervals (CI) were derived using t-values from scipy.stats, with 95% CI calculated accordingly.

Chip-to-chip normalization was conducted by calculating the mean relative affinity of wild-type RBD across all chips. Measured values were divided by this mean to address variability, and deviations from wild-type affinity were centered by subtracting 1 from normalized values.

To account for chip-to-chip variability, Imdevimab measurements were repeated on a different day with a new chip, showing consistent qualitative binding patterns, though intensity ratios were higher in the second experiment. Normalization to wild-type fluorescence ensured reliable inter-chip comparisons, yielding a strong correlation (R^2^ = 0.89, Supplementary Fig. 4). Detailed measurements and fitting plots are provided in the Supplementary Fig. 6–11).

The final dataset contains measured affinity values of triple mutation variants. For synergistic effect analysis, these measured values were transformed as follows:

1. Definition of Relative Affinity The relative affinity for a variant V with three mutations is defined as:

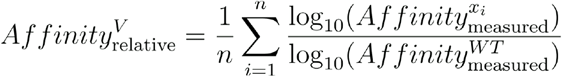

where *n* is the number of repetitions for the same variant, *x*_*i*_ is the measured affinity for each repetition, and *WT* represents the wild-type.
2. Mean Relative Affinity of Single Mutations The mean relative affinity for a single mutation *x* is calculated as the average relative affinity across all variants containing the mutation *x*. It is defined as:

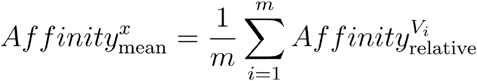

where *m* is the number of variants containing mutation *x, V*_*i*_ represents a variant containing mutation *x*.
3. The mean relative affinity of a double mutation pair *D* is defined as the average relative affinity of all variants that contain both mutations in *D*:

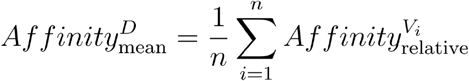

where *n* is the total number of variants containing both mutations in *D*, and *V*_*i*_ represents a variant containing the double mutation *D*.
4. Fold Change for Double Mutation Combination: The fold change quantifies the affinity-altering effect when mutations appear in combination. It is defined as:

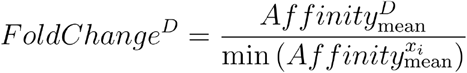

where *x*_*i*_ represents the single mutations within *D*. for triple mutation variant *V*:

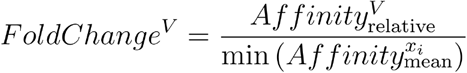

where *x*_*i*_ represents the single mutations within *V*.

### Sequence embedding and model training

To encode the receptor-binding domain (RBD) sequences, we employed three different embedding strategies:

1) Mutational Encoding: Each sequence was represented as a binary vector, where each possible mutation was assigned a value of 1 (present) or 0 (absent), relative to the wild-type RBD sequence. 2) One-Hot Encoding: Each amino acid was represented as a categorical one-hot vector, preserving full sequence information. 3) ESM2 Embeddings: Sequences were processed using esm2_t33_650M_UR50D, a pretrained protein language model. We extracted the representation from layer 15 to serve as the sequence embedding.

We benchmarked eight regression models for predicting relative affinity values using 5-fold cross-validation. 1) Gaussian Process (GP) Regression 2) Multi-Layer Perceptron (MLP) 3) Ridge Regression 4) Elastic Net 5) Random Forest (RF) 6)Gradient-boosted decision trees (XGBoost) 7) Support Vector Regression with linear (SVR_L) and RBF (SVR_rbf) kernel 8) ProteinNPT, a non-parametric transformer-based model with MSA_transformer embedding ^20^.

Performance was assessed using Pearson and Spearman correlation coefficients, mean squared error (MSE), and R^2^, providing a comprehensive evaluation of predictive accuracy across different model architectures.

### Figures and graphs

The RBD structures in Figure 1A and B were downloaded from PDB number 7B3O and the figures were generated using Pymol (Schroedinger, v.2). The binding interfaces for Bamlanivimab and Imdevimab are marked as previously cited ^5^. The binding interface for DBR3_03 was derived from PDB number 7ZRV by downloading the Cryo-EM structure of DBR3_03 bound to omicron ^15^. Subsequently, the binding interface on the RBD was determined by selecting all residues that were within 2.7 Å from DBR3_03. In Python, we used pandas, numpy, scipy.stats for data handling and statistical calculations, matplotlib, seaborn for figure generation. In R, we used the dplyr, ggplot2, tidyr, readr, iand graph for data wrangling and visualization. All plots were exported in PDF or PNG format and edited and assembled with Adobe Illustrator 2025.

## Code and data availability

https://github.com/LSSI-ETH/RBD_affinity

## Author Contribution

L.G. performed experiments. L.G. and S.J.M. designed experiments. L.G., J.T. and F.A. developed the platform. L.G. and MA.GC developed the variant assembly method. F.A. and L.G. fabricated the microfluidic molds. J.H. and S.R.T. designed and performed the epistasis and machine learning analysis with input from E.B.I and N.C. L.G., J.H., S.T.R., and S.J.M processed the data and wrote the manuscript.

## Acknowledgements

The authors would like to thank Simon Lietar for help and constant support with the automated MITOMI setup, and Gregoire Michielin for his help with image acquisition and processing. We thank Nikolas von den Eichen for his help with the liquid handling system, and Edgar Engel for fruitful discussions and help with data analysis. We would also like to thank Anthony Marchand and Bruno Correia for providing us with DBR3_03 and related discussions, as well as former and current LBNC members for helpful discussions and feedback. J.T. was supported by a grant from the Swiss National Science Foundation (Marie-Heim Voegtlin Fellowship, PMPDP3_171383). This work was also supported by the European Research Council under the European Union’s Horizon 2020 research and innovation program grant 723106 (to S.J.M.), a Swiss National Science Foundation NRP (National Research Program) 78 COVID-19 grant 198412 (to S.J.M.), and a Swiss National Science Foundation grant 182019 (to S.J.M.). Parts of Figure 1 were generated with the help of BioRender for: BioRender. Maerkl, S. (2025) https://BioRender.com/3214wb5, https://BioRender.com/kv4tdnw, https://BioRender.com/ptnxmnu

