## Supplementary Information for "High-throughput cell-free profiling of SARS-CoV-2 RBD variants enables rapid and quantitative in vitro affinity landscape mapping"

#### Supplementary Figures

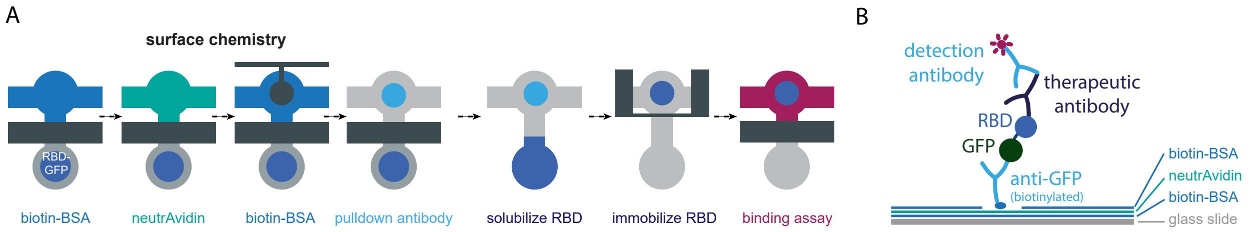


Supplementary Figure 1. (A) Schematic representation of the surface functionalization on the chip. All steps are performed with a closed neck valve, isolating the spotting chamber from the reaction chamber. First, the reaction chambers are coated with biotin-BSA. Subsequently, the surface is functionalized with neutrAvidin. The button valves are actuated to physically block the button area, and the remaining surfaces are passivized with another layer of biotin-BSA. Subsequently, the biotinylated anti-GFP pulldown antibody is flown across the chip and binds to the neutrAvidin-exposing button area. The RBDs are then solubilized and the unit chambers isolated by actuating the sandwich valves. The RBD-GFP fusion proteins are immobilized on the anti-GFP pulldown antibody, and the binding assay is performed. (B) Schematic representation of the binding assay. A layer of biotin-BSA is deposited on the glass slide, followed by a layer of neutrAvidin. The second layer of biotin-BSA is restricted to the area that is not covered by the button valve. The biotinylated anti-GFP antibody can then bind to the button region, and immobilizes the GFP-RBD in this region. Subsequently, therapeutic antibody or binder is flown, followed by the fluorescent detection antibody.

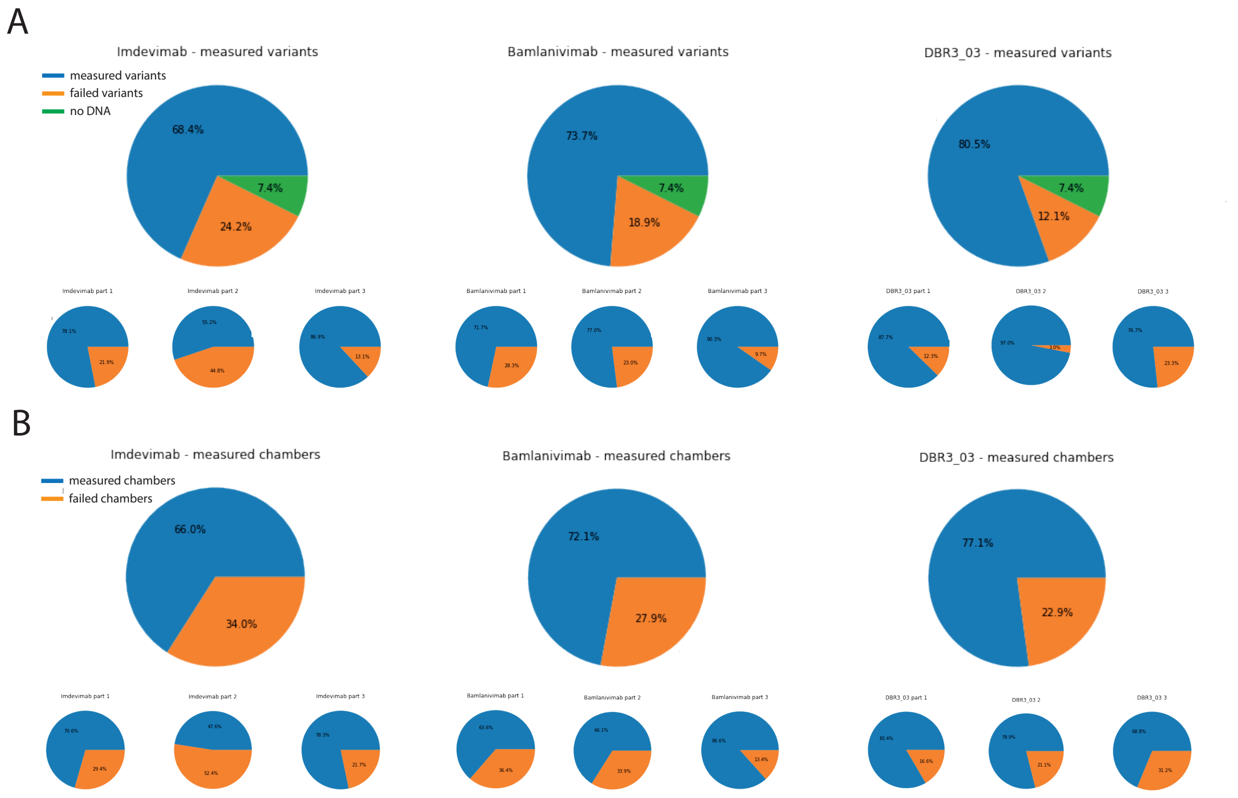


Supplementary Figure 2. Statistical analysis about the fraction of successfully analyzed variants and chambers. (A) Fractions of variants analyzed versus variants where no DNA was generated and variants with failed analysis for Imdevimab, Bamlanivimab and DBR3_03, as well as the ratios of successful vs non-successful variants on each chip for each variant. (B) Fractions of successful chambers vs non-successful chambers for Imdevimab, Bamlanivimab, and DBR3_03, as well as the respective fractions for each chip.

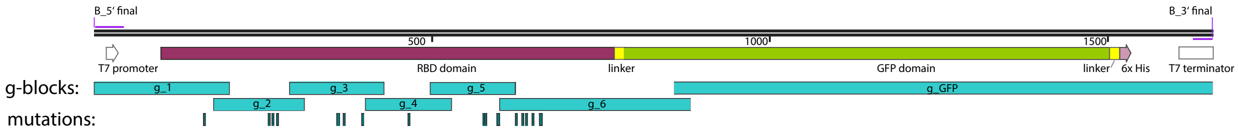


Supplementary Figure 3. Schematic map of the RBD-GFP fusion construct. The map contains the RBD (purple) and GFP (lime) features and the seven g-blocks (blue), as well as the distribution of the mutations (teal). Common features, including linkers, T7 promoter and terminator regions, and the primer binding sites are displayed, too.

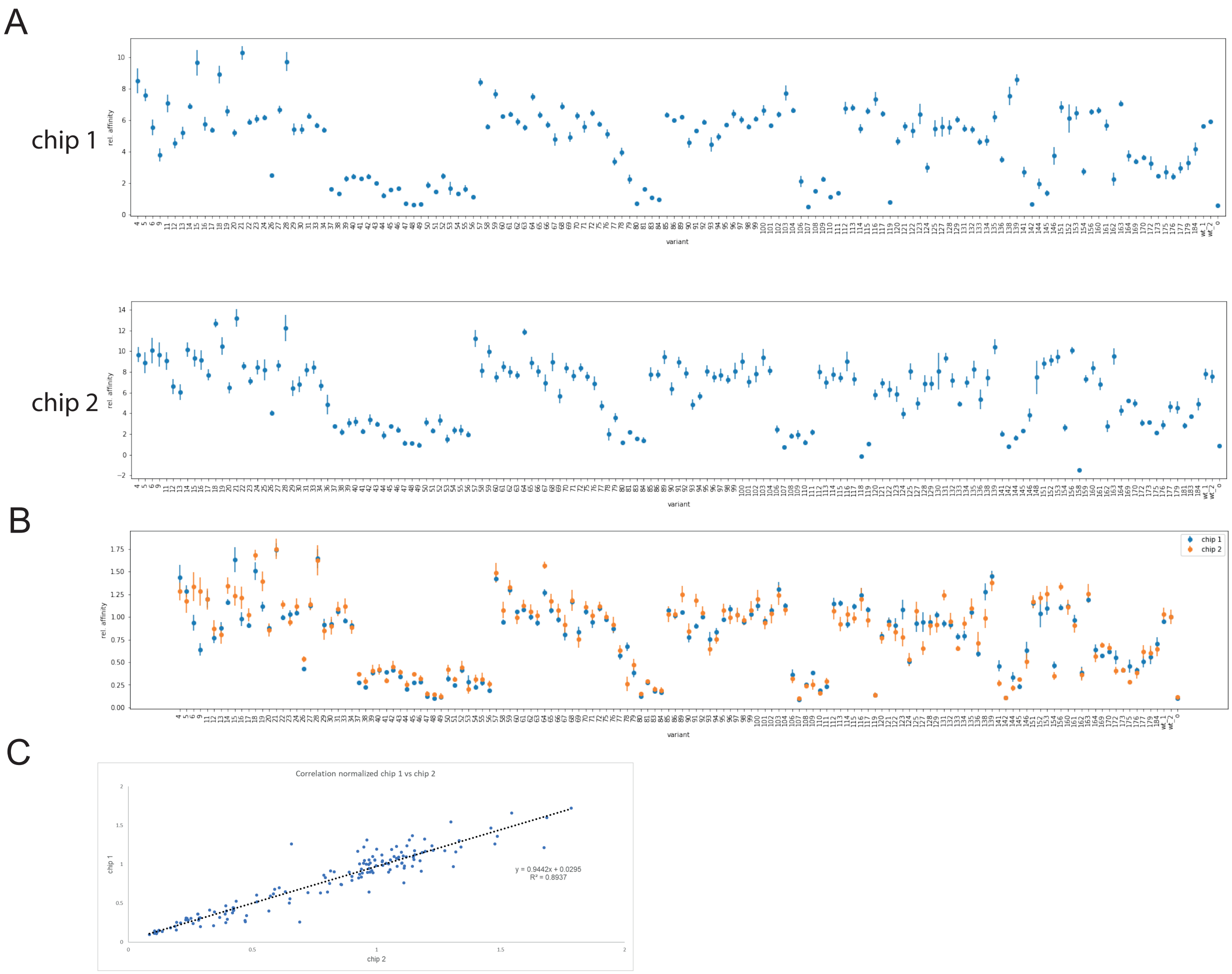


Supplementary Figure 4. Overview of the measured relative affinities of variants in part 1 against Imdevimab, to estimate the chip-to-chip variability. Despite the relative fluorescence for variants in chip 2 being higher, the qualitative differences between variations in binding affinities are comparable between the two chips.

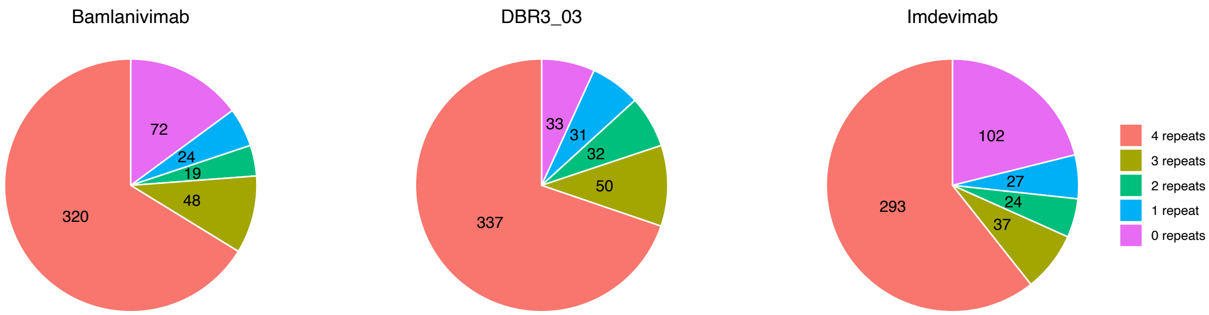


Supplementary Figure 5. Number of successful repeats performed for each variant in three binder datasets.

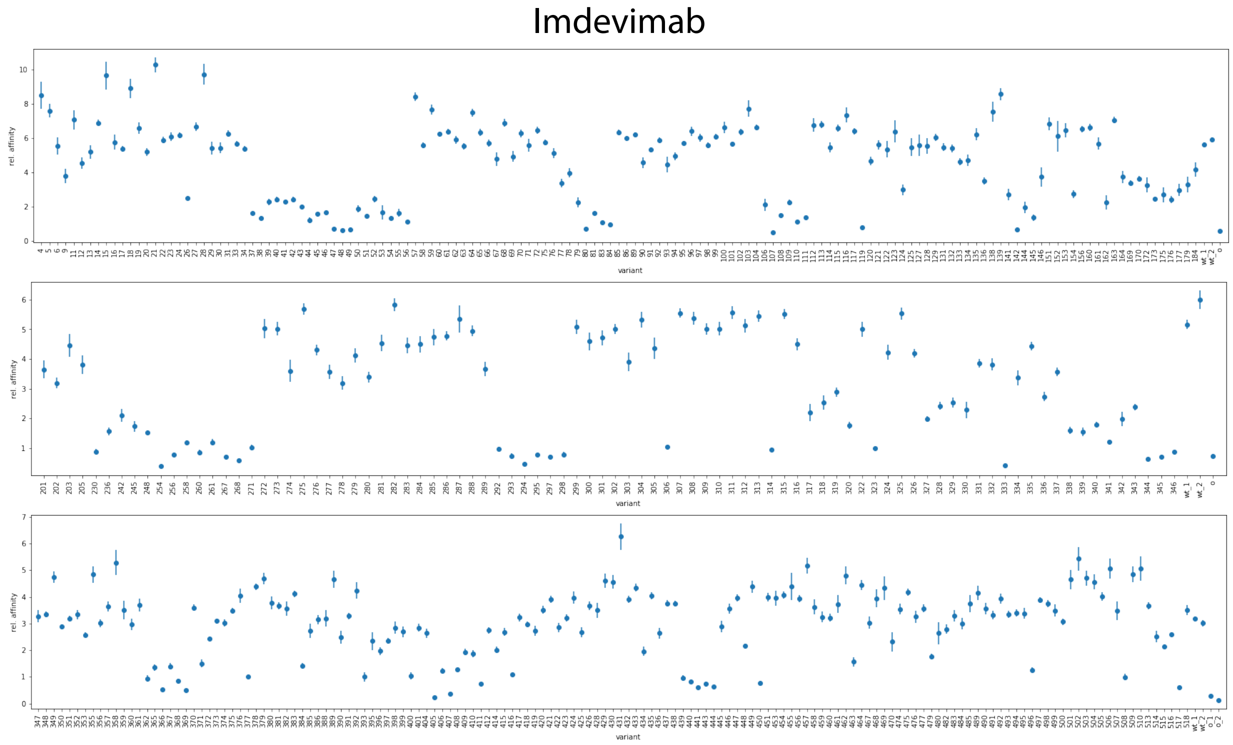


Supplementary Figure 6. Overview of relative affinities of all variants against Imdevimab across three chips.

#### Imdevimab

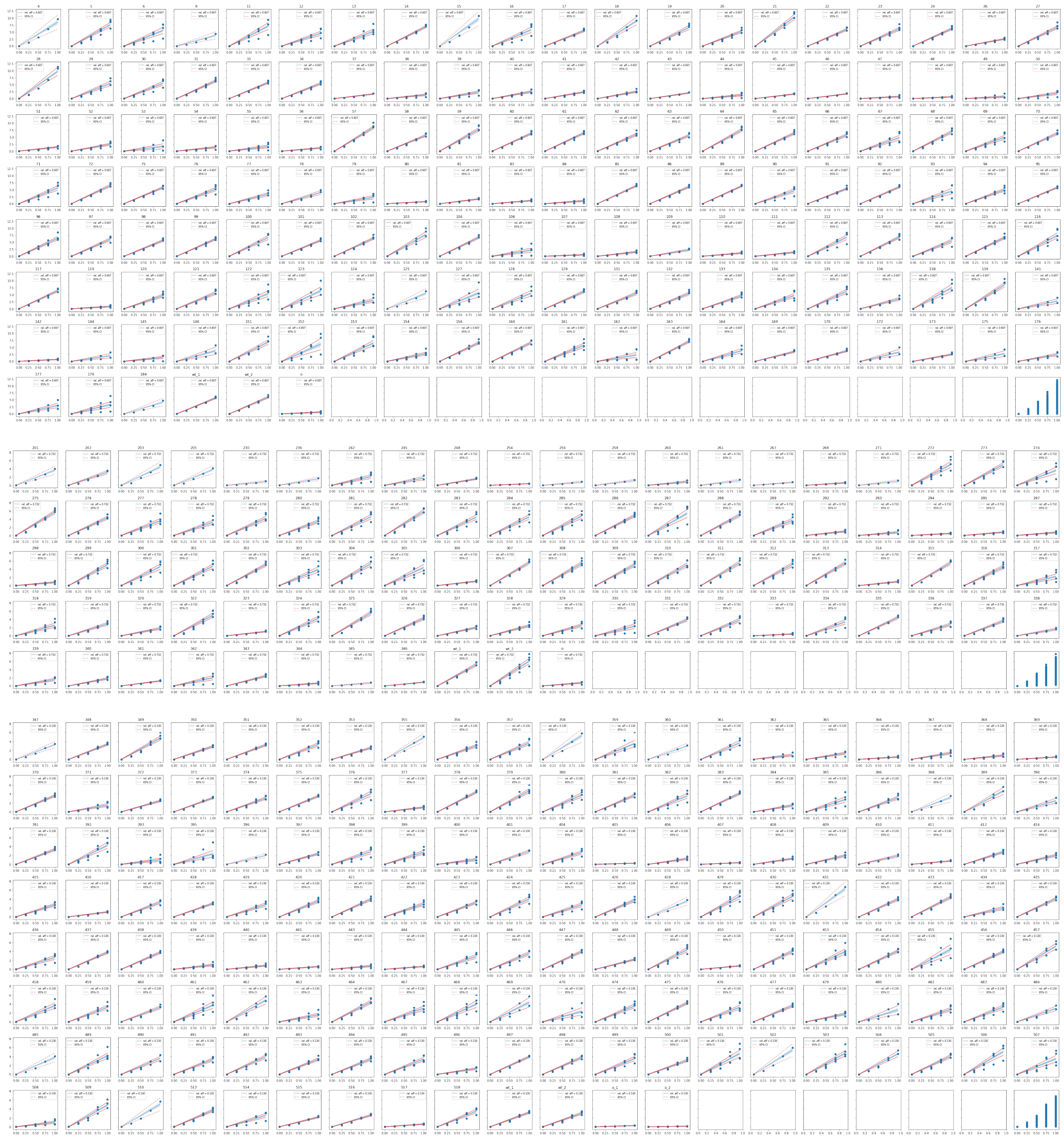

Supplementary Figure 7. Fits for Imdevimab including data points (blue), global linear regression (blue), standard error of slope (blue, dotted) and 95% confidence interval (red).

Bamlanivimab

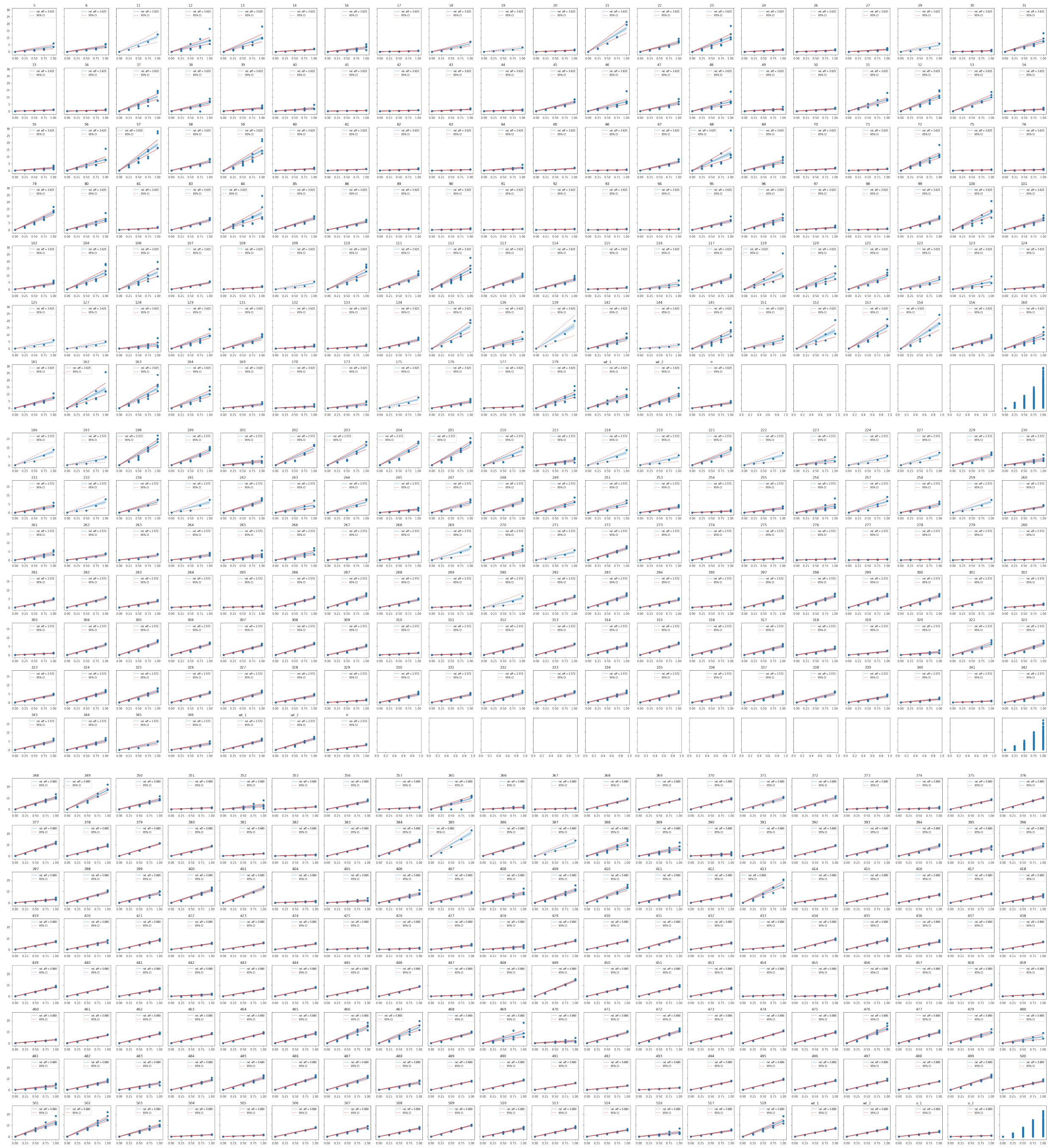

Supplementary Figure 8. Fits for Bamlanivimab including data points (blue), global linear regression (blue), standard error of slope (blue, dotted) and 95% confidence interval (red).

### Bamlanivimab

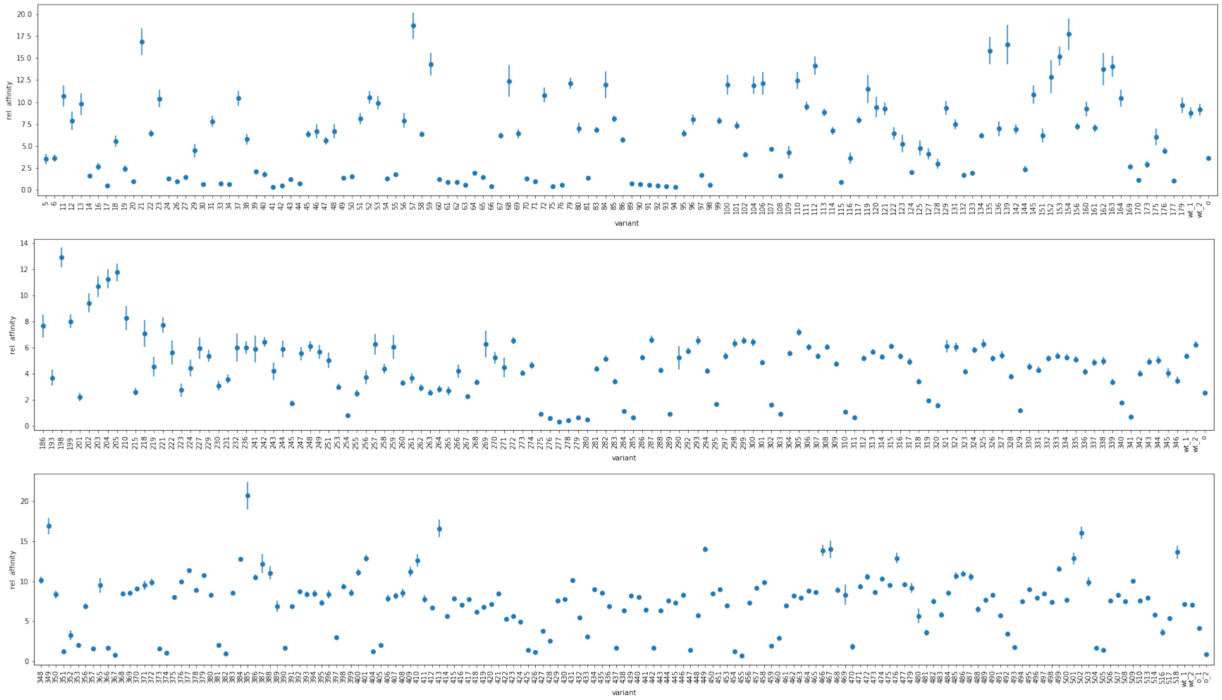

Supplementary Figure 9. Overview of relative affinities of all variants against Bamlanivimab across three chips.

### DBR3_03

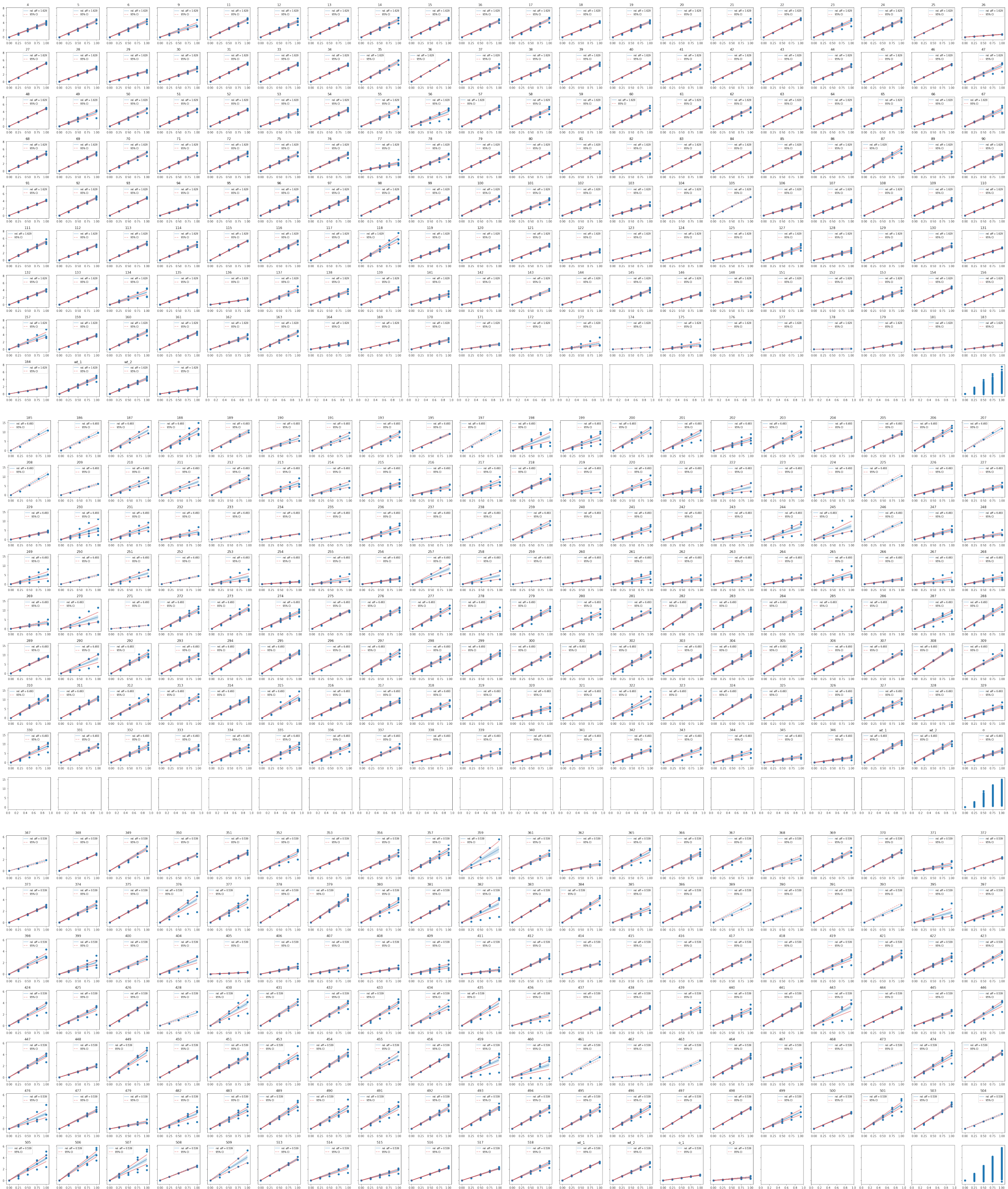

Supplementary Figure 10. Fits for DBR3_03 including data points (blue), global linear regression (blue), standard error of slope (blue, dotted) and 95% confidence interval (red).

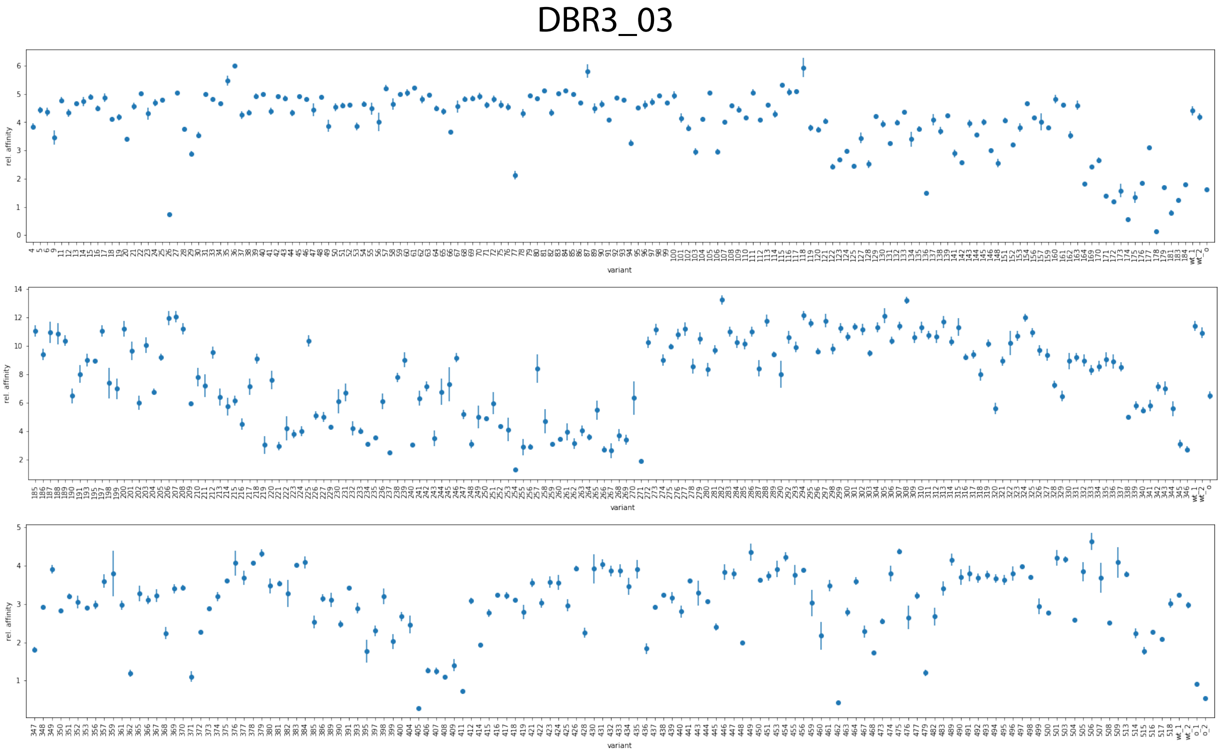


Supplementary Figure 11. Overview of relative affinities of all variants against DBR3_03 across three chips.

#### DNA used in this work

##### Primer sequences

The primers used in this work were:

Table 1: Primers used during this work

| | |
|---|---|
| B_5' final | GATCTTAAGGCTAGAGTACTAATACGACTCACTATAGGGAGACC |
| B_3' final | CAAAAAACCCCTCAAGACCCGTTTAGAG |

##### Touchdown PCR and PCR

Table 2: Taq touchdown PCR steps

| step | temperature [°C] | time [sec] |
|---|---|---|
| Initial denaturation | 95 | 30 |
| 10 cycles | 95<br>65, 64, 63, 62, 61, 60, 59, 58, 57, 56<br>68 | 30<br>30<br>100 |
| Final Extension | 68 | 5 min |

Table 3: Taq PCR steps

| step | temperature [°C] | time [sec] |
|---|---|---|
| Initial denaturation | 95 | 30 |
| 35 cycles | 95<br>57<br>68 | 30<br>30<br>100 |
| Final Extension | 68 | 5 min |

##### Exemplary sequence for Omicron

gatcttaaggctagagtacTAATACGACTCACTATAGGGAGACCACAACGGTTTCCCTCTAGAAATAATTTTGTTTAACTTAAGAAGGAGGAAAAAAAAATG
AGGGTTCAACCCACAGAATCAATAGTAAGATTCCCGAACATCACCAACCTGTGTCCGTTCGATGAGGTGTTTAATGCGACCCGCTTTGCATCCGT
CTACGCGTGGAATCGCAAACGTATCAGCAACTGCGTGGCCGATTATAGCGTACTGTATAACCTGGCTCCGTTCTTTACGTTTAAGTGCTACGGCG
TTAGCCCGACCAAATTGAATGATCTGTGCTTTACCAACGTGTACGCGGACAGCTTCGTTATCCGCGGTGACGAAGTTCGTCAGATTGCGCCAGGT
CAGACCGGTAACATCGCCGACTACAATTACAAGCTGCCGGACGACTTTACGGGCTGCGTCATCGCGTGGAATTCAAACAAACTGGATAGCAAGGT
GAGCGGCAACTACAACTACCTGTACCGCCTGTTTCGTAAGTCGAATCTGAAGCCGTTTGAGCGCGATATTTCCACGGAAATTTATCAAGCGGGTA
ATAAACCGTGTAATGGTGTGGCGGGCTTCAACTGCTACTTCCCGCTGCGTAGCTATAGCTTTCGTCCGACGTATGGCGTTGGTCATCAACCGTAT
CGTGTTGTGGTGTTAAGCTTCGAGTTGCTGCATGCACCGGCAACCGTTTGTGGTCCTAAAAAATCCACCAACCTGGTTAAGAACAAATGCGTTAA
CTTCGGCAGCGGTTCAGGCATGGTTAGCAAAGGTGAGGAACTGTTTACCGGCGTCGTGCCGATTTTGGTGGAGTTGGATGGCGACGTGAATGGT
CATAAATTCTCCGTTCGTGGTGAGGGCGAGGGCGACGCAACTAACGGCAAATTGACCCTGAAGTTCATCTGTACCACGGGTAAGTTGCCGGTTCC
GTGGCCGACTTTGGTGACCACTTTGACCTATGGCGTCCTTTGCTTCTCTCGTTATCCGGATCACATGAAACGTCACGATTTCTTTAAGAGCGCTA
TGCCGGAAGGCTATGTTCAAGAACGCACCATTTCCTTCAAGGACGACGGCACCTACAAAACCCGTGCGGAAGTCAAATTCGAGGGAGACACCCTG
GTGAACCGTATTGAGCTGAAGGGGATCGATTTTAAAGAAGACGGTAACATCTTAGGTCATAAGTTGGAGTATAACTTTAACAGCCATAATGTTTA
TATCACCGCTGATAAACAAAAAAATGGCATCAAGGCGTATTTCAAGATCCGTCACAACGTCGAGGACGGCAGCGTGCAGCTGGCGGACCACTACC
AACAGAATACCCCGATTGGTGACGGTCCGGTGTTACTCCCAGACAATCATTACCTGTCTACGCAGAGCGTGCTGAGCAAGGATCCGAACGAAAAA
CGTGATCACATGGTTCTGCTGGAAGATGTTACCGCGGCCGGTATTACCCATGGTATGGATGAGCTGTACAAGGGTTCGGGTAGTGGGCACCACCA
CCACCACCATTAATGAGATCCGGCTGCTAACAAAGCCCGAAAGGAAGCTGAGTTGGCTGCTGCCACCGCTGAGCAATAACTAGCATAACCCCTTG
GGGCCTCTAAACGGGTCTTGAGGGGTTTTTTG

#### Exemplary sequence for Wild Type

gatcttaaggctagagtacTAATACGACTCACTATAGGGAGACCACAACGGTTTCCCTCTAGAAATAATTTTGTTTAACTTAAGAAGGAGGAAAAAAAAATG
CGTGTACAGCCAACCGAATCAATTGTACGCTTTCCCAATATAACCAATCTATGCCCTTTCGGGGAAGTCTTCAATGCCACTCGTTTCGCTTCTGTG
TATGCATGGAACCGCAAGCGTATTTCCAACTGCGTTGCGGATTACTCCGTGTTGTATAATAGCGCGAGTTTCTCTACGTTTAAATGTTATGGTGT
GTCACCGACAAAACTGAACGATCTGTGCTTCACCAACGTGTACGCGGATAGCTTTGTTATTCGCGGCGACGAAGTTCGCCAGATCGCGCCAGGCC
AGACAGGCAAAATTGCCGATTATAACTATAAATTGCCGGACGATTTCACCGGATGCGTGATCGCGTGGAATAGCAACAACCTTGATAGCAAGGTC
GGTGGCAATTACAACTATCTCTACCGTCTGTTCCGCAAGAGCAATCTGAAACCCTTTGAGCGGGACATTTCGACTGAAATCTATCAAGCCGGGTC
GACGCCGTGTAACGGCGTGGAAGGATTTAACTGTTATTTTCCGCTGCAGTCGTACGGTTTTCAACCTACCAATGGCGTGGGCTACCAGCCGTACC
GAGTCGTGGTATTAAGTTTTGAGTTACTGCACGCCCCGGCAACGGTTTGTGGTCCGAAAAAAAGCACCAATCTTGTTAAAAATAAATGCGTCAAC
TTTGGTTCCGGGTCTGGTATGGTTTCCAAAGGAGAAGAACTGTTTACCGGTGTTGTACCAATTCTCGTAGAACTCGATGGAGATGTAAACGGGCA
TAAATTTTCAGTGCGCGGCGAGGGCGAAGGAGATGCCACAAACGGCAAACTGACCCTTAAATTTATTTGCACGACCGGCAAATTACCAGTTCCTT
GGCCTACGCTGGTCACCACGCTCACCTATGGGGTATTATGCTTTAGCCGCTATCCGGATCACATGAAACGCCATGATTTCTTTAAAAGTGCTATG
CCAGAAGGTTATGTACAGGAACGCACGATTAGCTTTAAAGATGATGGGACGTATAAAACCCGCGCCGAGGTAAAATTTGAAGGAGATACCTTAGT
AAACCGCATTGAACTCAAGGGGATTGATTTTAAAGAGGACGGTAACATTCTGGGTCATAAACTTGAGTACAACTTTAACTCACACAACGTTTACA
TTACCGCGGATAAACAGAAGAACGGTATTAAAGCGTACTTTAAGATTCGCCATAACGTCGAAGATGGCAGTGTTCAGCTGGCCGATCATTATCAG
CAGAACACGCCGATTGGCGATGGCCCTGTTTTGTTACCGGATAACCATTATTTATCGACTCAGAGCGTCTTAAGTAAAGATCCAAACGAGAAACG
CGATCACATGGTTCTCTTAGAAGATGTTACCGCCGCCGGCATTACACATGGCATGGATGAACTGTATAAAGGTTCCGGGTCTGGTCATCATCATC
ACCATCACtgaTGAGATCCGGCTGCTAACAAAGCCCGAAAGGAAGCTGAGTTGGCTGCTGCCACCGCTGAGCAATAACTAGCATAACCCCTTGGGG
CCTCTAAACGGGTCTTGAGGGGTTTTTTG

Supplementary Table 1. Model performance in predicting binding affinity with Bamlanivimab

| | Mutational Encoding | | | | One-Hot Encoding | | | | ESM2 Representation | | | |
|---|---|---|---|---|---|---|---|---|---|---|---|---|
| **Model** | MSE | Pearson | Spearman | $R^2$ | MSE | Pearson | Spearman | $R^2$ | MSE | Pearson | Spearman | $R^2$ |
| GP | 0.11±0.03 | 0.77±0.07 | 0.8±0.04 | 0.59±0.1 | 0.11±0.03 | 0.77±0.07 | 0.8±0.04 | 0.59±0.1 | 0.11±0.04 | 0.77±0.07 | 0.81±0.03 | 0.58±0.13 |
| LR | 0.11±0.03 | 0.76±0.06 | 0.82±0.03 | 0.58±0.09 | - | - | - | - | - | - | - | - |
| MLP | 0.13±0.08 | 0.74±0.14 | 0.79±0.09 | 0.5±0.27 | 0.12±0.03 | 0.77±0.06 | 0.81±0.05 | 0.54±0.1 | 0.12±0.03 | 0.75±0.05 | 0.79±0.03 | 0.54±0.09 |
| Ridge | 0.11±0.03 | 0.76±0.06 | 0.82±0.03 | 0.57±0.08 | 0.11±0.03 | 0.76±0.06 | 0.82±0.03 | 0.58±0.08 | 0.11±0.03 | 0.78±0.06 | 0.81±0.04 | 0.59±0.11 |
| ElasticNet | 0.11±0.03 | 0.77±0.05 | 0.82±0.03 | 0.58±0.07 | 0.12±0.04 | 0.76±0.06 | 0.82±0.03 | 0.56±0.12 | 0.11±0.03 | 0.78±0.06 | 0.81±0.04 | 0.6±0.09 |
| RF | 0.09±0.03 | 0.81±0.07 | 0.83±0.05 | 0.65±0.11 | 0.09±0.03 | 0.81±0.07 | 0.83±0.05 | 0.65±0.11 | 0.13±0.02 | 0.74±0.04 | 0.77±0.03 | 0.52±0.04 |
| XGBoost | 0.12±0.03 | 0.8±0.08 | 0.81±0.06 | 0.56±0.1 | 0.12±0.03 | 0.8±0.08 | 0.81±0.06 | 0.56±0.1 | 0.14±0.02 | 0.68±0.04 | 0.7±0.03 | 0.45±0.06 |
| SVM_rbf | 0.11±0.03 | 0.78±0.07 | 0.8±0.04 | 0.58±0.11 | 0.1±0.03 | 0.79±0.07 | 0.82±0.04 | 0.62±0.11 | 0.11±0.03 | 0.78±0.05 | 0.81±0.04 | 0.6±0.09 |
| SVM_linear | 0.14±0.02 | 0.76±0.05 | 0.81±0.03 | 0.47±0.05 | 0.12±0.03 | 0.76±0.06 | 0.81±0.03 | 0.54±0.07 | 0.1±0.03 | 0.79±0.05 | 0.81±0.04 | 0.61±0.09 |
| ProteinNPT | | | | | | | | | 0.44±0.15 | 0.76±0.07 | 0.79±0.05 | 0.56±0.12 |
| ProteinNPT multi-targets | | | | | | | | | 0.38 | 0.81 | 0.82 | 0.61 |

Supplementary Table 2. Model performance in predicting binding affinity with Imdevimab

| | Mutational Encoding | | | | One-Hot Encoding | | | | ESM2 Representation | | | |
|---|---|---|---|---|---|---|---|---|---|---|---|---|
| **Model** | MSE | Pearson | Spearman | $R^2$ | MSE | Pearson | Spearman | $R^2$ | MSE | Pearson | Spearman | $R^2$ |
| GP | 0.06±0.02 | 0.81±0.08 | 0.81±0.08 | 0.66±0.13 | 0.06±0.02 | 0.81±0.08 | 0.81±0.08 | 0.66±0.13 | 0.05±0.01 | 0.84±0.03 | 0.83±0.03 | 0.71±0.05 |
| LR | 0.06±0.01 | 0.81±0.06 | 0.82±0.04 | 0.66±0.1 | - | - | - | - | - | - | - | - |
| MLP | 0.12±0.15 | 0.69±0.34 | 0.75±0.2 | 0.23±1 | 0.06±0.02 | 0.8±0.09 | 0.8±0.08 | 0.62±0.16 | 0.09±0.04 | 0.77±0.15 | 0.75±0.14 | 0.45±0.31 |
| Ridge | 0.06±0.01 | 0.81±0.06 | 0.82±0.04 | 0.66±0.1 | 0.06±0.01 | 0.81±0.06 | 0.82±0.04 | 0.66±0.1 | 0.13±0.16 | 0.7±0.33 | 0.77±0.16 | 0.22±1.03 |
| ElasticNet | 0.05±0.01 | 0.84±0.03 | 0.84±0.02 | 0.69±0.05 | 0.05±0.01 | 0.84±0.03 | 0.83±0.02 | 0.69±0.05 | 0.08±0.06 | 0.75±0.21 | 0.77±0.14 | 0.53±0.41 |
| RF | 0.05±0.02 | 0.83±0.08 | 0.83±0.06 | 0.69±0.13 | 0.05±0.02 | 0.83±0.08 | 0.83±0.06 | 0.69±0.13 | 0.06±0.01 | 0.8±0.06 | 0.79±0.04 | 0.63±0.09 |
| XGBoost | 0.06±0.03 | 0.81±0.13 | 0.79±0.15 | 0.65±0.22 | 0.06±0.03 | 0.81±0.13 | 0.79±0.15 | 0.65±0.22 | 0.07±0.01 | 0.77±0.06 | 0.76±0.04 | 0.59±0.09 |
| SVM_rbf | 0.06±0.01 | 0.82±0.06 | 0.83±0.05 | 0.68±0.1 | 0.06±0.01 | 0.82±0.06 | 0.83±0.04 | 0.67±0.1 | 0.05±0.01 | 0.85±0.03 | 0.84±0.03 | 0.72±0.05 |
| SVM_linear | 0.06±0.01 | 0.82±0.06 | 0.82±0.03 | 0.67±0.1 | 0.06±0.01 | 0.82±0.06 | 0.82±0.03 | 0.66±0.1 | 0.05±0.02 | 0.83±0.09 | 0.82±0.07 | 0.69±0.14 |
| ProteinNPT | | | | | | | | | 0.37±0.09 | 0.79±0.08 | 0.76±0.08 | 0.61±0.15 |
| ProteinNPT multi-targets | | | | | | | | | 0.45 | 0.68 | 0.68 | 0.35 |

Supplementary Table 3. Model performance in predicting binding affinity with DBR3_03

| | Mutational Encoding | | | | One-Hot Encoding | | | | ESM2 Representation | | | |
|---|---|---|---|---|---|---|---|---|---|---|---|---|
| **Model** | MSE | Pearson | Spearman | $R^2$ | MSE | Pearson | Spearman | $R^2$ | MSE | Pearson | Spearman | $R^2$ |
| GP | 0.04±0.01 | 0.71±0.06 | 0.68±0.06 | 0.5±0.09 | 0.04±0.01 | 0.71±0.06 | 0.68±0.06 | 0.5±0.09 | 0.04±0.01 | 0.75±0.05 | 0.73±0.04 | 0.53±0.09 |
| LR | 0.04±0.01 | 0.74±0.06 | 0.72±0.06 | 0.53±0.09 | - | - | - | - | - | - | - | - |
| MLP | 0.08±0.07 | 0.61±0.24 | 0.65±0.09 | 0.12±0.78 | 0.04±0.01 | 0.73±0.07 | 0.7±0.06 | 0.5±0.1 | 0.13±0.08 | 0.73±0.05 | 0.73±0.03 | -0.41±0.92 |
| Ridge | 0.06±0.01 | 0.74±0.06 | 0.72±0.06 | 0.33±0.04 | 0.05±0.01 | 0.74±0.06 | 0.72±0.06 | 0.43±0.06 | 0.05±0.02 | 0.72±0.06 | 0.7±0.04 | 0.4±0.27 |
| ElasticNet | 0.04±0.01 | 0.75±0.06 | 0.72±0.06 | 0.55±0.09 | 0.04±0.01 | 0.75±0.06 | 0.72±0.06 | 0.55±0.09 | 0.04±0.01 | 0.74±0.05 | 0.72±0.04 | 0.49±0.14 |
| RF | 0.04±0.01 | 0.75±0.06 | 0.72±0.06 | 0.55±0.09 | 0.04±0.01 | 0.75±0.06 | 0.72±0.06 | 0.55±0.09 | 0.04±0.01 | 0.73±0.05 | 0.71±0.04 | 0.53±0.06 |
| XGBoost | 0.04±0.01 | 0.75±0.05 | 0.72±0.05 | 0.54±0.07 | 0.04±0.01 | 0.75±0.05 | 0.72±0.05 | 0.54±0.07 | 0.05±0.01 | 0.69±0.07 | 0.66±0.07 | 0.46±0.1 |
| SVM_rbf | 0.04±0.01 | 0.75±0.06 | 0.73±0.07 | 0.55±0.1 | 0.08±0.01 | 0.72±0.06 | 0.71±0.07 | 0.08±0.05 | 0.04±0.01 | 0.75±0.05 | 0.74±0.05 | 0.55±0.08 |
| SVM_linear | 0.04±0.01 | 0.74±0.06 | 0.73±0.06 | 0.53±0.1 | 0.04±0.01 | 0.74±0.06 | 0.73±0.07 | 0.53±0.1 | 0.06±0.03 | 0.74±0.07 | 0.73±0.05 | 0.37±0.39 |
| ProteinNPT | | | | | | | | | 0.58±0.05 | 0.68±0.05 | 0.65±0.07 | 0.41±0.08 |
| ProteinNPT multi-targets | | | | | | | | | 0.53 | 0.74 | 0.72 | 0.50 |

**ProteinNPT model used MSA transformer embedding
