## Supplementary material for "High-throughput cell-free profiling of SARS-CoV-2 RBD variants enables rapid and quantitative in vitro affinity landscape mapping": supplementary_data_annotation.rtf

Supplementary Data – “triple_mutant_synergy.csv”
This comma-separated file lists every RBD variant that carries exactly three point mutations and was assayed for binding affinity against each of the three antibodies in the study. The table is already filtered to include only variants for which all three single-mutation affinities are available, so that additive and synergistic effects can be recomputed directly from the file.
Antibody: string – antibody tested (Bamlanivimab, Imdevimab, or DBR3)
Mutations: string – three space-separated mutation codes (e.g., E484A S375F N440K) listed in ascending position order
Mean_Affinity_WT_3mut: numeric – mean relative affinity of the triple mutant, normalised to wild-type (values < 1 indicate reduced affinity)
Individual_Score_1: numeric – mean relative affinity of the first single mutation (matches first code in Mutations)
Individual_Score_2: numeric – mean relative affinity of the second single mutation
Individual_Score_3: numeric – mean relative affinity of the third single mutation
fold_change: numeric – synergy metric calculated as Mean_Affinity_WT_3mut divided by the minimum of the three Individual_Score values (values < 1 denote stronger-than-additive escape)
Rank: integer – rank of the triple mutant within its antibody dataset after sorting by ascending fold_change (1 = strongest synergistic escape)
All affinity values are normalised (unit-less). Wild-type affinity = 1 by definition.
Supplementary Data – “g_block_list.csv”
name: name of the g block
sequence: sequence of the g block
mutations: mutations in the g block, “wt” means wild type, “o” denotes for Omicron variant
